# Phenylalanine Treatment Suppresses Necrotrophic and Biotrophic Pathogens in Mono- and Dicotyledonous Plants

**DOI:** 10.64898/2026.07.26.740754

**Authors:** Yigal Elad, Dalia Rav-David, Michal Oren-Shamir

**Affiliations:** Department of Plant Pathology and Weed Research, Institute of Crop Protection, Volcani Institute, Agricultural Research Organization, Volcani Institute, Rishon LeZion 7534509, Israel; Department of Ornamental Plants and Agricultural Biotechnology, Institute of Plant Sciences, Volcani Institute, Agricultural Research Organization, Volcani Institute, Rishon LeZion 7534509, Israel

**Keywords:** Phenylalanine, fungal pathogens, Oomycetes pathogens, bacterial diseases, necrotrophy, biotroph, dicot, monocot

## Abstract

L-phenylalanine (Phe) is converted via the phenylpropanoid pathway into phenolic compounds, and elevated phenolic/flavonoid levels are generally associated with enhanced plant resistance. We previously showed that exogenous Phe suppresses the necrotrophic fungus *Botrytis cinerea* in petunia, chrysanthemum, determinant tomato, and several postharvest diseases. Here, we expanded evaluation to a broad range of pathosystems, including an indeterminate greenhouse tomato cultivar, multiple dicot species, and the monocot wheat. Phe, applied as spray or drench across concentrations, was consistently effective at ≥4 mM. It reduced disease severity in diverse systems: *Sclerotinia sclerotiorum* (tomato, sweet basil, cucumber, lettuce), *Leveillula taurica* and *Oidium neolycopersici* (tomato), *Podosphaera xanthii* (cucumber), wheat foliar pathogens (*Blumeria graminis* f. sp. *tritici*, *Zymoseptoria tritici*, *Puccinia triticina*, *P. striiformis* f. sp. *tritici*), the oomycetes *Pseudoperonospora cubensis* (cucumber leaves) and *Pythium aphanidermatum* (roots), bacterial pathogens (*Pseudomonas syringae* pv. *tomato* and *Clavibacter michiganensis* subsp. *michiganensis*) and Tomato brown rugose fruit (ToBRFV) that belongs to the *Tobamovirus* genus. Phe was effective on both young and mature leaves and often performed comparably to chemical fungicides; combinations rarely improved control, except for enhanced activity with pyrimethanil against *B. cinerea* in tomato. Synergistic effects were observed when Phe was combined with an adjuvant against tomato powdery mildews. A formulated product (NaturaFend 550 SP) was more effective than non-formulated Phe in *B. cinerea* (tomato) and *P. xanthii* (cucumber). Application timing also influenced efficacy, with treatment 5 days before infection providing superior control compared with 0, 3, or 7 days. Overall, Phe effectively controlled biotrophic and necrotrophic fungi, oomycetes, bacterial pathogens and a virus across diverse crops under experimental greenhouses and commercial like conditions.

## 1 Introduction

The phenylpropanoid pathway (PPP) converts L-phenylalanine into diverse phenolic compounds, including lignin, flavonoids, anthocyanins, and coumarins (Treutter 2006). Many of these metabolites possess antioxidant and antifungal activities (Sudheeran et al. 2020; Treutter 2006) and contribute to pigment production, UV protection, and defense against biotic and abiotic stresses (Sharma et al. 2019; Yadav et al. 2020; Yao et al. 2021). The PPP is central to induced plant defense, and higher phenolic or flavonoid levels are associated with increased pathogen resistance (Conrath et al. 2002; Dixon et al. 2002; Sivankalyani et al. 2016). Activation of this pathway can prime plant defenses (Conrath et al. 2002) and enhancing flavonoid content through breeding or metabolic engineering may further improve resistance (Forkmann & Martens 2001).

Plants efficiently uptake and metabolize L-phenylalanine into phenylpropanoids (Kumari et al. review). Exogenous application of L-phenylalanine induces defense responses, as shown by reduced *Botrytis cinerea* infection in petunia and tomato (Oliva et al. 2020) and decreased postharvest diseases such as anthracnose (*Colletotrichum gloeosporioides*), stem-end rot (*Lasiodiplodia theobromae*), and green mold (*Penicillium digitatum*) in fruits (Kumar Patel et al. 2020). This effect appears indirect, as L-phenylalanine does not inhibit pathogen germination but instead enhances host defenses, including increased phenylalanine ammonia-lyase (PAL) activity and reduced *Erysiphe pisi* conidia germination (Bahadur et al. 2012).

Following our previous findings that L-phenylalanine application suppresses *B. cinerea* in petunia and tomato (Oliva et al. 2020), chrysanthemum (Kumar Varun et al. 2020), and postharvest diseases in mango, avocado, and citrus (Kumar Patel et al. 2020), we extended our study to assess its effects on a broader range of pathogens, including necrotrophic and biotrophic fungi, oomycetes, and bacteria in both dicot and monocot plants.

The present study deals with plant pathogens of variable nature. *B. cinerea* (causal agent of gray mold) and *S. sclerotiorum* (causes white mold) are both necrotrophs that infect high number of plant hosts including many agricultural crops (Elad et al. 2016; Hegedus & Rimmer 2005). Two powdery mildew pathogens infect tomato: *Leveillula taurica*, a broad host-range species that colonizes the full leaf tissue, and *Oidium neolycopersici*, a tomato-specific pathogen that develops superficially on upper leaf surfaces (Elad et al. 2009; Jacob et al. 2008). In cucurbits, *Podosphaera xanthii* causes powdery mildew in cucumber (Elad et al. 2021a; Reuveni et al. 1993). All three are obligate biotrophic fungi. In wheat, we examined foliar diseases caused by obligate biotrophs, including *Blumeria graminis* f. sp. *tritici* (powdery mildew), *Puccinia triticina* (leaf rust), and *Puccinia striiformis* f. sp. *tritici* (stripe rust), as well as the hemi-biotrophic fungus *Zymoseptoria tritici*, the causal agent of Septoria tritici blotch (Couture, 1980; Lipps Madden 1989). In cucumber, two oomycetes were evaluated: *Pseudoperonospora cubensis*, an obligate biotroph causing downy mildew, and *Pythium aphanidermatum*, a necrotroph responsible for damping-off (Barnea et al. 2022; Philosoph et al. 2019; Sivan et al. 1984). Two hemibiotrophic bacterial pathogens of tomato were studied: the Gram-negative *Pseudomonas syringae* pv. *tomato*, which causes bacterial speck, and the Gram-positive *Clavibacter michiganensis* subsp. *michiganensis*, the causal agent of bacterial canker (Louws et al. 2001; Verma et al. 2024). Finaly, the virus Tomato brown rugose fruit (ToBRFV) that belongs to the *Tobamovirus* genus. The virus causes symptoms including mosaic and distortion of leaves and brown, wrinkly spots (rugose) on fruits and it spreads short distances by mechanical contact (Luria et al. 2017; Salem et al. 2016, 2023).

Building on our previous studies demonstrating control of postharvest diseases and *Botrytis cinerea* in tomato, petunia, and chrysanthemum using L-phenylalanine (Kumar Patel et al. 2020; Kumar Varun et al. 2020; Oliva et al. 2020), this study aimed to evaluate its efficacy across a wide range of pathosystems. In tomato, we extended the analysis to an indeterminate greenhouse cultivar, examining different L-phenylalanine concentrations, spatial effects across leaf ages, comparisons and combinations with botryticides, and optimized application timing using formulated treatments.

Beyond tomato, we assessed its activity against the necrotroph *Sclerotinia sclerotiorum* in multiple plant species, powdery mildews of tomato and cucumber, and fungal pathogens of wheat (a monocot), including biotrophs and hemi-biotrophs. We also evaluated its effects on two oomycete pathogens (airborne and soilborne), two bacterial diseases of tomato and the virus ToBRFV in tomato.

## 2 Materials and methods

### 2.1 Growth conditions

Plants were grown in a 20-mesh net house (April–October 2018–2024) or in a glasshouse with 25% shade netting and whitewash (Livnat, Tapazol, Israel) during the rest of the year. Day/night temperatures ranged from 18–32°C. Plants were maintained pest- and pathogen-free until treatments were applied. Plants were grown in 1–5 L pots containing a peat–tuff (≤8 mm) mixture (7:3, v/v). Fertigation was applied 2–3 times daily via drip irrigation using 5:3:8 NPK fertilizer (120:30:150 mg L ¹ N:P:K; EC 2.2 dS m ¹), maintaining 25–50% drainage (Elad et al. 2010). All experiments were conducted at the Volcani Institute, Rishon LeZion, Israel (31°58′09″N, 34°48′02″E).

### 2.2 Plant treatments

Plants were sprayed or drenched with L-phenylalanine (P), P formulations, or fungicides. Spray applications were performed using a 1 L hand sprayer to ensure full canopy coverage with fine droplets, applying 8–15 mL per plant depending on size. Drench applications consisted of 10–30 mL per pot, covering the root zone with ∼5% drainage.

P was applied mainly as an aqueous solution of L-phenylalanine (Merck-Millipore, USA), and in some experiments as a formulated product, NaturaFend 550 SP (550 g kg ¹ L-phenylalanine; ICA International Chemicals, South Africa). P was used at 1–32 mM depending on the experiment.

Fungicides and elicitors included: Switch (cyprodinil + fludioxonil, Syngenta), Mythos/Scala SC (pyrimethanil, Bayer), Topaz 200 EW (penconazole, Syngenta), Heliosulfur (sulfur, Action Pin), Dynon (propamocarb HCl, Bayer), Bion 50 WG (acibenzolar-S-methyl, Syngenta), monopotassium phosphate (MKP; KH PO, ICL Haifa), and the adjuvant Shatah 90 (alkyl phenol oxide, Adama/Makhteshim, Israel).

### 2.3 Plants

Tomato (*Solanum lycopersicum*) cv. Ikram (Syngenta, Israel) seedlings were obtained from Hishtil Nurseries (Israel) 4–6 weeks after sowing, transplanted to pots, and grown for 4–6 additional weeks before pathogen inoculation.

Cucumber (*Cucumis sativus*) cv. Manny (Genesis Seeds, Israel) was grown in peat–tuff medium. To prevent damping-off by *Pythium* spp. in non-*Pythium* experiments, the medium was drenched once at sowing with 0.25% Dynon (propamocarb HCl, Bayer). Cucumber plants were used as: (i) seedlings for *Pythium aphanidermatum* assays (5 seedlings per pot), (ii) 5–7-week-old plants (1 L pots, one plant per pot) for foliar disease studies, and (iii) long-term plants in 5 L pots (one plant per container).

Wheat (*Triticum aestivum*) was grown in 3 L pots (8 seeds per pot) for 7–13 weeks. Cv. Mabruk was used as an inoculum source for foliar pathogens, while cv. Mabruk and cv. Bet Hashita were used as experimental hosts.

Lettuce (*Lactuca sativa*) cv. Raviv was obtained at ∼1 month of age and transplanted into 1 L pots.

Sweet basil (*Ocimum basilicum*) cv. Peri seedlings were raised in a commercial nursery and transplanted at 3–4 weeks of age. Plugs containing 3–5 plants were grown in 1 L pots (one plug per pot), and each plug was treated as one plant.

### 2.4 Pathogens and diseases

*Botrytis cinerea* isolate BcI16 (Elad, 1989) was grown on PDA at 22°C. Conidia from 14-day-old cultures were suspended in 0.1% glucose + KH PO and adjusted to 5 × 10 conidia/mL. Ten µL drops were used to inoculate tomato leaflets (typically middle leaves, L5–L6; in some experiments also L9–L10 were inoculated). Plants were incubated in a humid chamber at 22–24°C with 14 h light. Disease severity was scored as lesion area (0–100%). In some assays, 3 mm mycelial discs from 3–4-day-old cultures were placed on leaflets, and lesion diameter/area was measured.

*Sclerotinia sclerotiorum* Sc1 (Elad, 2000) was grown on half-strength PDA at 22°C. Three-mm mycelial discs from 4-day-old cultures were placed on leaves of tomato, sweet basil, cucumber, and lettuce. Plants were incubated under humid conditions at 22–24°C. Lesion diameter was measured daily and converted to rot area.

Two tomato powdery mildew pathogens were used: *Leveillula taurica* (LtPM) and *Oidium neolycopersici* (OnPM). Infected tomato plants served as inoculum sources. Conidia were washed from symptomatic leaves, suspended in water (10 conidia/mL), microscopically verified, and sprayed onto experimental plants. Disease severity was scored weekly as leaf coverage (0–100%).

Cucurbit powdery mildew (*Podosphaera xanthii*, PxPM) was obtained from naturally infected plants. Conidia were collected in water (10 /mL) and sprayed onto cucumber plants. Severity was assessed weekly as percent leaf coverage.

Wheat foliar diseases were established using cv. Mabruk as an inoculum bridge. Field-infected plants showing symptoms of powdery mildew, leaf rust, stripe rust, and Septoria blotch were transferred to the greenhouse and placed among experimental pots (one inoculum pot per 10 pots). Natural spread was allowed. For *Zymoseptoria tritici*, sprinkler irrigation (10 min/day for 1 week) was used to enhance infection. Pathogens included *Blumeria graminis* f. sp. *tritici*, *Z. tritici*, *Puccinia triticina*, and *P. striiformis* f. sp. *tritici*. Disease severity (0–100%) was assessed on 10 mature leaves per pot.

Cucumber downy mildew (*Pseudoperonospora cubensis*) developed naturally in net-house conditions (Nov–Apr) and was scored as leaf coverage (0–100%).

Cucumber damping-off was caused by *Pythium aphanidermatum*, grown on PDA at 25°C. Mycelium was transferred to autoclaved millet, incubated for 5 days, homogenized, and mixed into the growth medium at 0.25% (w/w). Seedlings were transplanted into infested medium, and mortality was recorded every 2 days.

Tomato bacterial speck (*Pseudomonas syringae* pv. *tomato*) developed naturally in net-house experiments and was rated as leaf coverage (0–100%).

Bacterial canker (*Clavibacter michiganensis* subsp. *michiganensis*) was cultured on LB agar at 28°C, suspended in water (5 × 10 CFU/mL), and sprayed onto tomato plants. Plants were maintained at 25 ± 2°C, and disease severity was rated on a 0–100 scale.

An Israeli isolate of ToBRFV, ToBRFV-IL (KX619418.1) was used for Tomato brown rugose fruit virus ToBRFV artificial infection. Tomato plants were sprayed with carborundum powder suspension and the leaves were rubbed with a phosphate buffer solution (0.01 M, pH 7.0) supplemented with crushed leaves from the ToBRFV-infected tomato plants. The severity of ToBRFV symptoms were evaluated on a scale of 0-100% (Alon et al. 2021; Klap et al. 2020). RT-PCR analysis was carried out according to Alon et al. (2021).

### 2.5 Linear growth of pathogens in culture

Mycelial growth was assessed using 3 mm discs from the margins of 3-day-old cultures. *Rhizoctonia solani* was grown at 28°C on PDA, *Botrytis cinerea* and *Sclerotinia sclerotiorum* at 22°C, and *Pythium aphanidermatum* at 25°C. Cultures were grown on PDA amended with 1–16 mM P. Colony diameter was measured daily, colony area calculated, and growth rate determined as the change in area between selected time points.

### 2.7 Calculations and statistics

Rot area and leaf area were calculated from measured data. Percentage data were arcsine-transformed prior to analysis. Area under the disease progress curve (AUDPC) was calculated where applicable. Standard errors (SE) are shown in figures and tables. Replication ranged from 5 to 10 per treatment and replicates were randomly arranges in blocks (Tables 1–3).

**Table 1:**
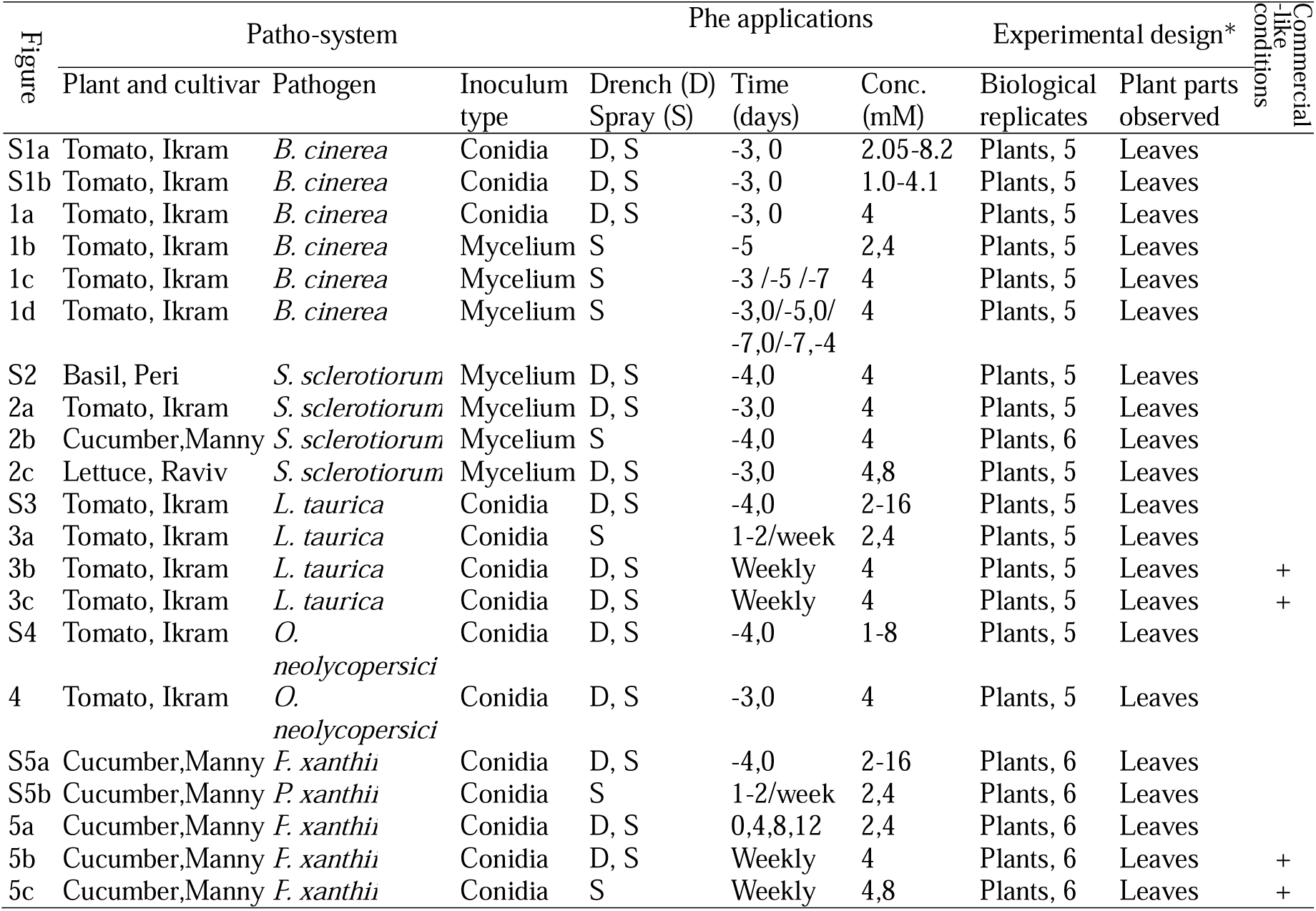
Summary of hosts, pathogens, Phe application and disease assessment described throughout the article for dicot plants infected by fungal pathogens.

**Table 2:**
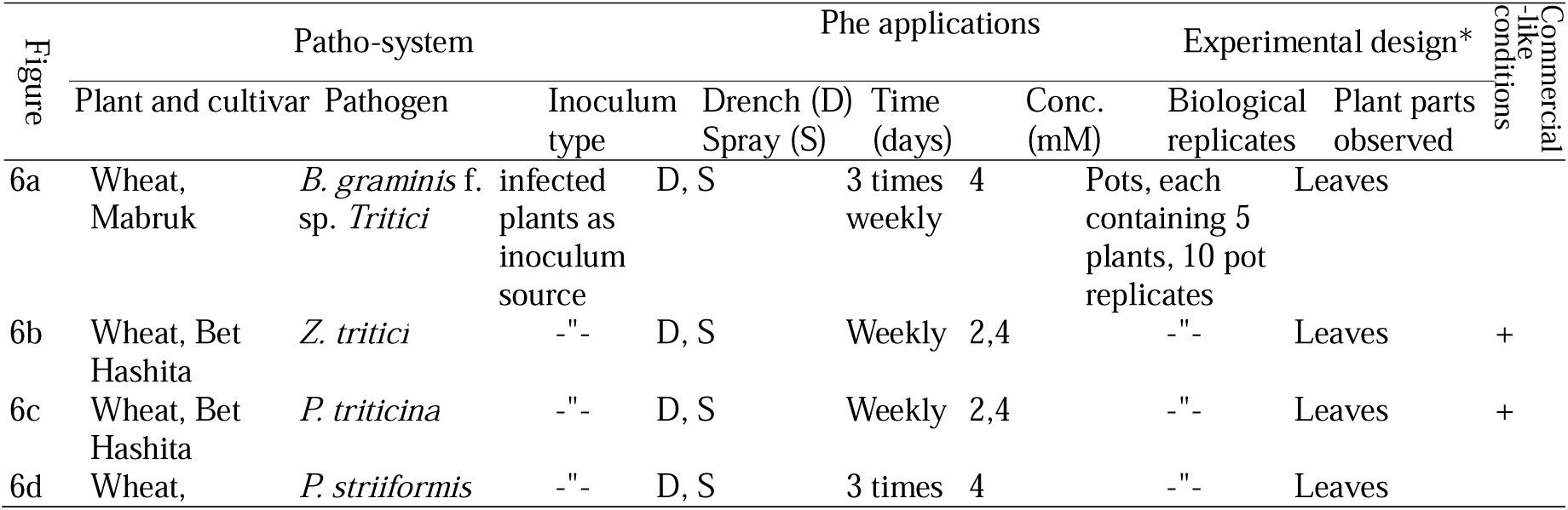
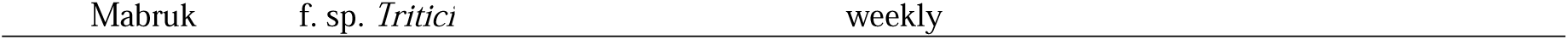
Summary of pathogens, Phe application and disease assessment described throughout the article for wheat plants.

**Table 3:** Summary of hosts, pathogens, Phe application and disease assessment described throughout the article for Oomycetes, bacteria and a virus.

| Figure | Patho-system |  | Phe applications |  |  |  | Experimental design* |  | Commercial-like conditions |
| --- | --- | --- | --- | --- | --- | --- | --- | --- | --- |
|  | Plant and cultivar | Pathogen | Inoculum type | Drench (D) Spray (S) | Time (days) | Conc. (mM) | Biological replicates | Plant parts observed |  |
| 7 | Cucumber, Manny | *P. cubensis* | Natural | D, S | Weekly | 4 | Plants, 6 | Leaves | + |
| S6 | Cucumber, Manny | *P. aphanidermatum* | Milet seed inoculum | S | Pre-transplanting | 4 | Pots | Seedling |  |
| 8 | Cucumber, Manny | *P. aphanidermatum* | -"- | D, S | 3, 6, and 10 after sowing | 2, 4 | Pots, each containing 5 plants, 8 pot replicates | Seedling |  |
| 9a | Tomato, Ikram | *P. syringae* pv. *tomato* | Natural | D, S | -4, 0 | 4 | Plants, 5 | Leaves |  |
| 9b | Tomato, Ikram | *P. syringae* pv. *tomato* | Natural | S | -3, 0 | 4-12 | Plants, 5 | Leaves |  |
| 9c | Tomato, Ikram | *C. michiganensis* subsp. *michiganensis* | Bacterial cells | D, S | -3, 0 | 4 | Plants, 5 | Leaves |  |
| 10ab | Tomato, Ikram | ToBRFV | Infected tomato tissue | D, S | 0,7,14 | 4 | Plants, 5 | Whole plant |  |

\* All experiments random blocks

Data (disease severity and AUDPC) were analyzed by ANOVA followed by Tukey–Kramer HSD test, using JMP software (SAS Institute, Cary, NC, USA; α = 0.05). Different letters indicate significant differences among treatments.

Disease reduction (DR, %) was calculated as:

DR = 100 − 100 × (Tn / Tcon), where Tn is the disease level in treated plants, and Tcon is the untreated control.

Synergism between treatments was evaluated using the Abbott-based model (Kosman & Cohen 1996). Expected disease reduction (DRexp) was calculated as:

DRexp = DR1 + DR2 − (DR1 × DR2 / 100).

Observed reduction (DRobs) was determined experimentally. Synergism factor (SF) = DRobs / DRexp; SF = 1 indicates additivity, and SF > 1 indicates synergy.

Tables 1-3 summarize the patho-systems that are described in the current article, application schedule, and assessment methods.

## 3 Results

### 3.1 Detailed study of Phe treatment effect on *B. cinerea* in greenhouse tomato cv. Ikram

Previous studies reported that Phe controls *B. cinerea* in processing tomato, petunia, and chrysanthemum (Oliva et al. 2019; Kumar Varun et al. 2020). Here, we evaluated spatial effects across leaf age, concentrations (≥1 mM), compatibility with fungicides, formulation, and application timing. Disease severity was higher on older (middle) leaves. Phe significantly reduced rot at 2.05–8.20 mM on older leaves via both spray and drench, while on younger leaves, reductions occurred mainly with 4.1–8.2 mM sprays and all drench treatments. Overall, drench applications were more effective than sprays, with efficacy increasing with concentration (Figure S1a).

Sprays of 1.0–4.1 mM performed similarly to 0.1% Switch (∼75% disease reduction), while drenches at 2.05–4.1 mM were superior to equivalent sprays (Figure S1b). Both application modes at 4 mM matched 0.06% Switch (∼89% reduction). Phe combined with Switch showed no added benefit over single treatments. Mythos (0.15%) reduced disease by 45%, and its combination with Phe drench showed additive effect (SF = 1.02), whereas no improvement was observed with spray combinations (Figure 1a).

**FIGURE 1.**
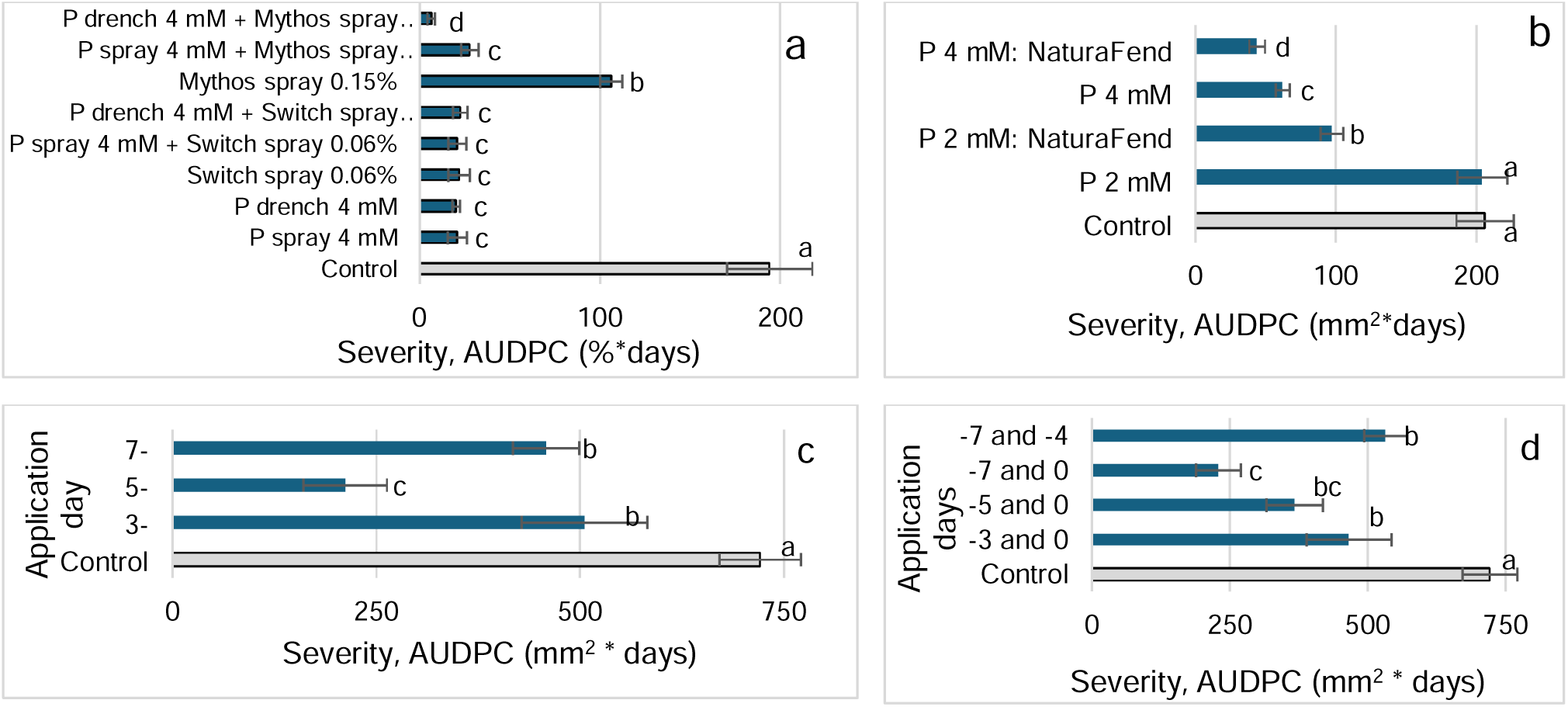
Control of *Botrytis cinerea* gray mold (BcGM) on tomato cv. Ikram by Phe, fungicides, and a Phe formulation (NaturaFend 550 SP). (a) Effect of Phe (4.0 mM) applied by spray or drench, Switch, Mythos (pyrimethanil, Bayer), and their combinations. Phe was applied at −3 and 0 days, and fungicides at 0 days before infection; disease was assessed up to 13 days post-infection. (b) Effect of non-formulated vs. formulated Phe (NaturaFend) at 2–4 mM applied by spray. Treatments were applied at −5 days before mycelial disc infection and assessed up to 4 days post-infection. (c) Effect of a single 4 mM Phe spray applied 3–7 days before mycelial disc infection. (d) Effect of two 4 mM Phe sprays applied 0–7 days before mycelial disc infection. BcGM severity after conidial infection (a) was rated on a 0–100% scale (0 = healthy; 100 = maximum rot). After mycelial disc infection, severity reflects lesion area (b,c,d). Area under the disease progress curve (AUDPC) was calculated over the assessment period. Different letters indicate significant differences (one-way ANOVA with Tukey–Kramer HSD, α = 0.05). Error bars represent SE.

Formulated Phe (NaturaFend) at 2–4 mM was more effective than non-formulated Phe (Figure 1b). A single 4 mM spray reduced disease by 30–36% when applied 3 or 7 days before infection, and by 71% when applied 5 days prior (Figure 1c). Two-spray programs (7 and 0 days before infection) provided the most consistent control (Figure 1d).

### 3.2 Phe controls white mold (*Sclerotinia sclerotiorum*) on tomato, sweet basil, cucumber, and lettuce leaves

*S. sclerotiorum* was evaluated as a second necrotrophic pathogen targeted by Phe (Figures 2 and S2). Both spray and drench applications reduced white mold (SsWM) across all tested crops. In tomato, 4 mM Phe reduced disease by 39% (drench) and 71% (spray) (Figure 2a), while in sweet basil, reductions were 42% and 66%, respectively (Figure S2). In cucumber, 4 mM spray reduced SsWM by 53%, though Topaz was significantly more effective (Figure 2b). In lettuce, disease reduction ranged from 36–52% with 4 mM drench or spray and 8 mM spray, whereas 8 mM drench provided the strongest control (84%) and was significantly superior to other treatments (Figure 2c).

**FIGURE 2.**
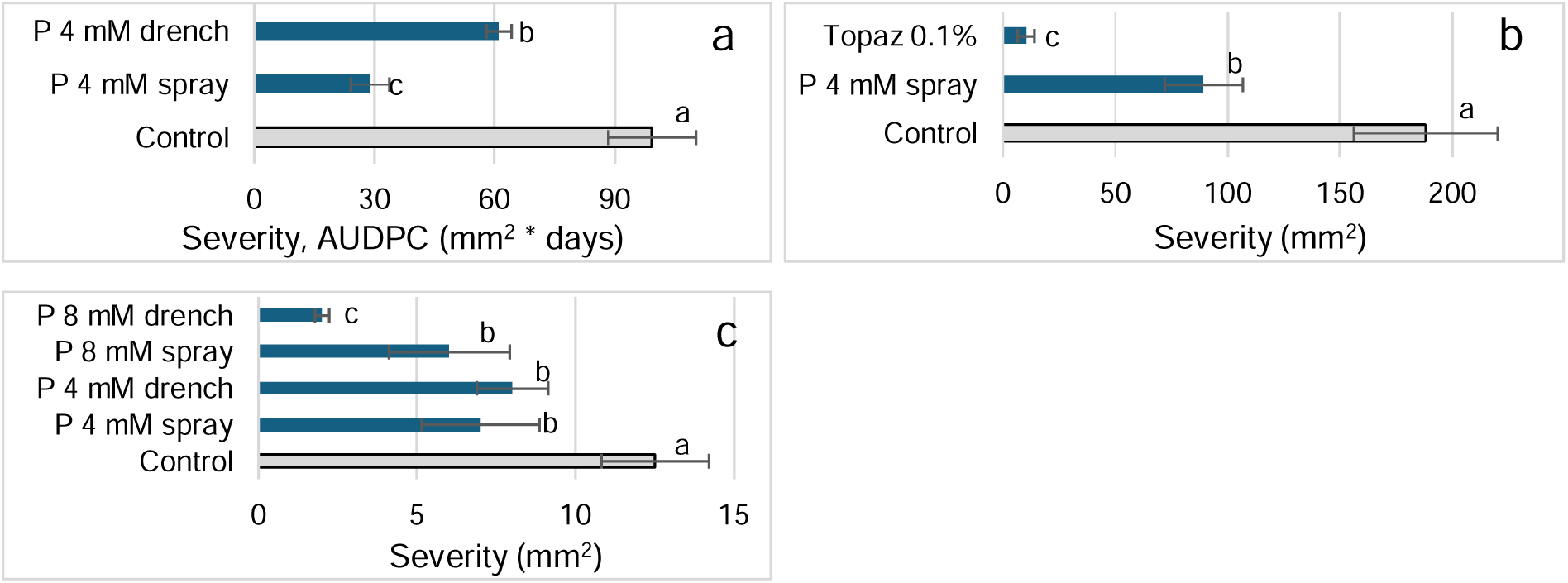
Effect of P on *Sclerotinia sclerotiorum* white rot (SsWM) caused by mycelial disc infection on tomato, sweet basil, cucumber, and lettuce leaves. (a) Tomato SsWM control by 4 mM P applied as spray or drench at −3 and 0 days before inoculation; disease assessed over 9 days. (b) Cucumber SsWM control by 4 mM P spray applied at −4 and 0 days, compared with Topaz (penconazole, Syngenta®); disease assessed at 3 days post-inoculation. (c) Lettuce SsWM control by 4- and 8-mM P spray applied at −3 and 0 days; disease assessed at 3 days post-inoculation. Disease severity was quantified by lesion diameter and converted to rot area (a–c). Area under the disease progress curve (AUDPC) was calculated for (a). Different letters indicate significant differences (one-way ANOVA with Tukey–Kramer HSD, α = 0.05). Error bars represent SE.

### 3.3 Control of two powdery mildew diseases of tomato by Phe treatment

#### 3.3.1 Application of Phe against *Leveillula taurica* powdery mildew (LtPM)

Phe applied as a spray or drench at 4 and 16 mM significantly reduced LtPM severity by 46–85%, whereas 2 mM showed no significant effect (Figure S3). Increasing 2 mM Phe applications from once to twice weekly did not improve control, while 4 mM spray remained consistently effective (Figure 3a).

**FIGURE 3.**
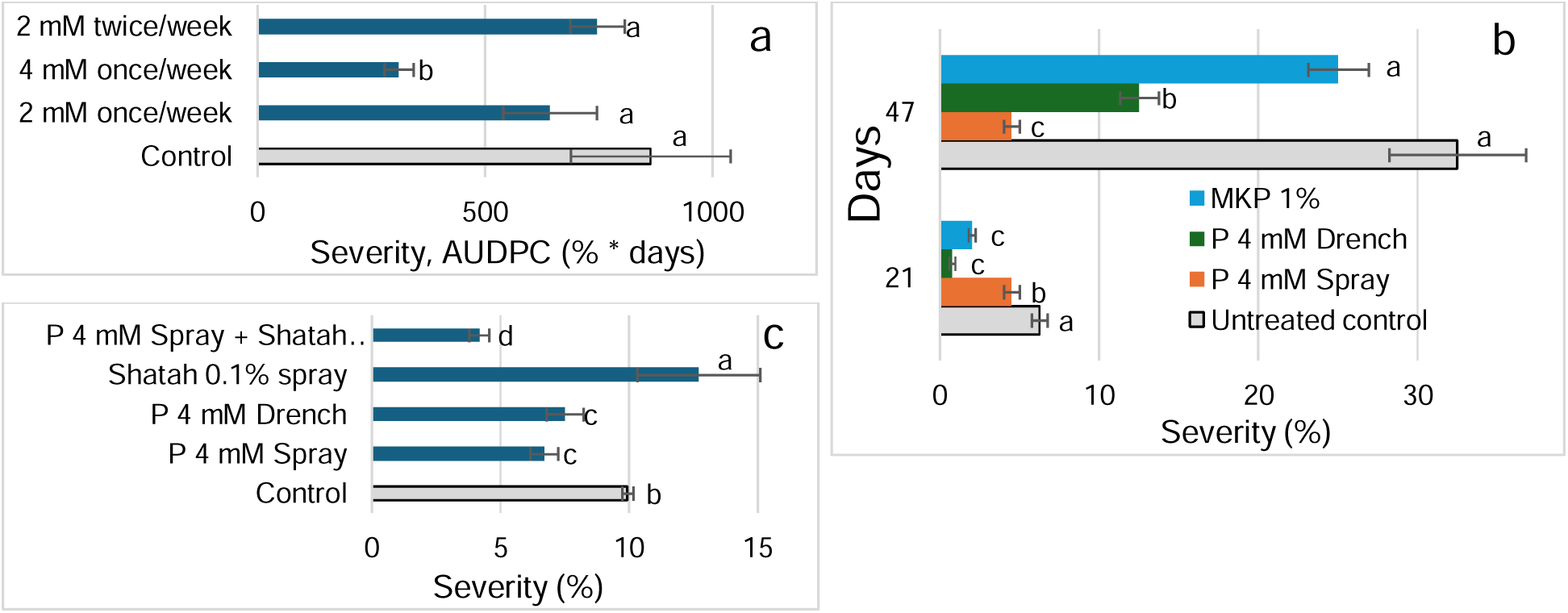
Effect of P on *Leveillula taurica* powdery mildew (LtPM) of tomato. (a) Effect of 2 mM P applied once or twice weekly and 4 mM P applied once weekly over 8 weeks; disease severity was assessed up to 73 days after treatment initiation. (b) Effect of 4 mM P (spray and drench) and monopotassium phosphate (MKP; KH PO) applied at 0, 7, 14, 21, 34, and 41 days; disease severity was assessed at 21 and 47 days after treatment initiation. (c) Effect of 4 mM P (spray or drench), Shatah 90 (alkyl phenol oxide 920 g/L, Adama/Makhteshim), and their combination applied at 0, 7, 14, and 21 days; disease severity was assessed at day 24. Disease severity was rated on a 0–100 scale (0 = healthy, 100 = fully covered leaves). AUDPC was calculated in (a). Different letters indicate significant differences (one-way ANOVA with Tukey–Kramer HSD, α = 0.05). Error bars represent SE.

The efficacy of 4 mM Phe was compared with monopotassium phosphate (MKP; KH PO), a known powdery mildew suppressor (Reuveni et al. 1993). Both spray and drench applications of 4 mM Phe significantly reduced LtPM, whereas MKP showed no effect under the tested conditions (Figure 3b).

Combining 4 mM Phe spray with the adjuvant Shatah 90 enhanced disease suppression relative to Phe alone, while the adjuvant alone had no effect (Figure 3c). The interaction was strongly synergistic (SF = 14.25).

#### 3.3.2 Effect of Phe on tomato *Oidium neolycopersici* powdery mildew (OnPM)

Phe was applied at 1–8 mM to tomato plants infected with *O. neolycopersici* (Figure S4). Foliar spray treatments at all concentrations were highly effective, achieving >91% disease suppression. In contrast, drench application showed a dose-dependent response: 1 mM had no significant effect, 2 mM reduced disease by 63%, and 4–8 mM resulted in complete suppression of OnPM (Figure S4). Combination of shatah 90 and P was significantly more effective than the Phe spray alone (Figure 4). The synergism between the shatah 90 and the Phe spray was significant (SF=1.21).

**FIGURE 4.**
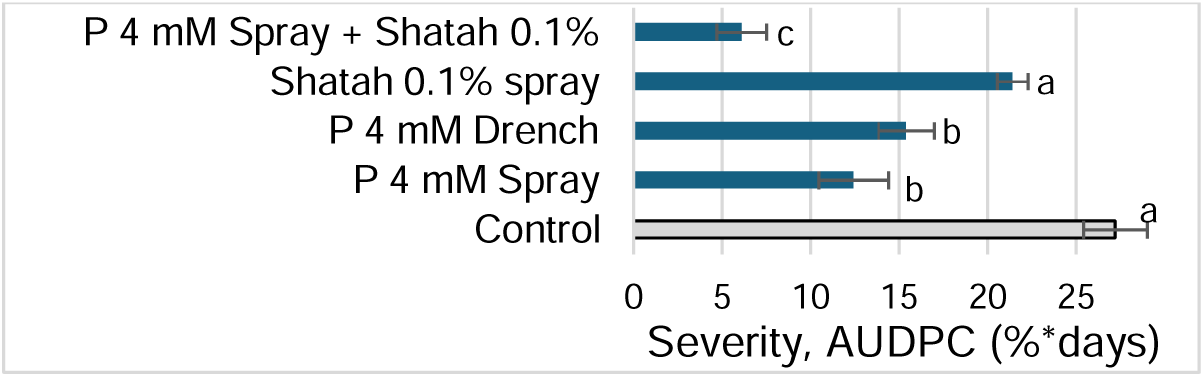
Effect of P on *Oidium neolycopersici* powdery mildew (OnPM) of tomato. Effect of 4 mM P (spray or drench), Shatah 90 (alkyl phenol oxide 920 g/L, Adama/Makhteshim), and their combination. Treatments were applied at −3 and 0 days before inoculation and then weekly until 28 days post-inoculation; disease was assessed up to day 35. Disease severity was rated on a 0–100 scale (0 = healthy, 100 = fully covered leaves). AUDPC was calculated in (b). Different letters indicate significant differences (one-way ANOVA with Tukey–Kramer HSD, α = 0.05). Error bars represent SE.

### 3.4 Effect of Phe on *Podosphaera xanthii* cucurbit powdery mildew (PxPM) on cucumber

Phe was evaluated against the biotrophic pathogen *P. xanthii* on cucumber (Figure 5 and S5). On seedlings, drench and spray applications of 2–4 mM Phe significantly reduced disease incidence, with 2 mM achieving 32–50% suppression and 4 mM providing greater control (Figure 5a). On plants with true leaves, 2–16 mM Phe applied in five treatments showed that 2 mM was ineffective, while 4–16 mM significantly reduced disease severity by 51–65% (Figure S5a). Increasing the frequency of 2 mM spray from once to twice weekly did not improve control (Figure S5b).

**FIGURE 5.**
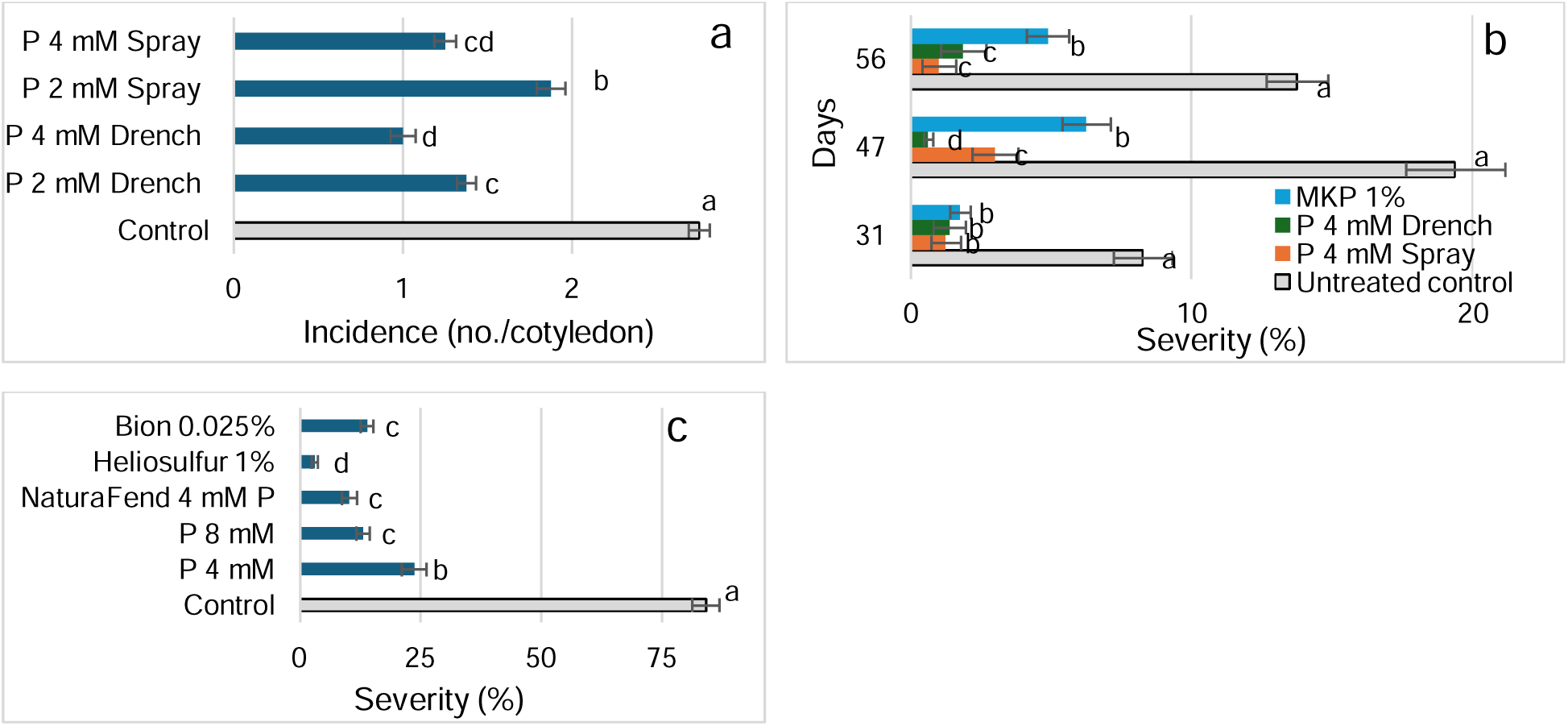
Effect of P on *Podosphaera xanthii* powdery mildew (PxPM) of cucumber. (a) Effect of P (2- and 4- mM) applied by drench or spray to germinating seedlings. Drench was applied once pre-emergence and twice post-emergence (4-day intervals), and spray twice post-emergence. Disease developed naturally (no artificial inoculation), and incidence was assessed at 16 days after seeding. (b) Effect of 4 mM P (spray and drench) and monopotassium phosphate (MKP; KH PO) applied weekly for 7 weeks; disease severity was assessed at 31, 47, and 56 days after treatment initiation. (c) Effect of weekly spray applications (7 total) of 4- and 8-mM P, NaturaFend 4 mM P, Heliosulfur 1.0% (sulfur a.i., Actionpin), and Bion 0.025% (acibenzolar-S-methyl, Syngenta); disease was assessed at 54 days after treatment initiation. Disease severity was rated on a 0–100 scale (0 = healthy, 100 = fully covered leaves). Different letters indicate significant differences (one-way ANOVA with Tukey– Kramer HSD, α = 0.05). Error bars represent SE.

Compared with monopotassium phosphate (MKP), 4 mM Phe provided significantly higher disease suppression (Figure 5b). Further comparisons showed that 4 mM Phe reduced PxPM by 72%, while 8 mM Phe, NaturaFend 550 SP, and Bion achieved ∼85% reduction; Heliosulfur was most effective with 96% suppression, outperforming all other treatments (Figure 5c).

### 3.5 Control of foliar diseases in wheat by Phe

Phe efficacy was further evaluated on foliar diseases of wheat, a monocotyledonous species (Figure 6). Wheat powdery mildew (*Blumeria graminis* f. sp. *tritici*) was reduced by 4 mM Phe applied as a spray or drench by 63 and 55%, respectively (Figure 6a).

**FIGURE 6.**
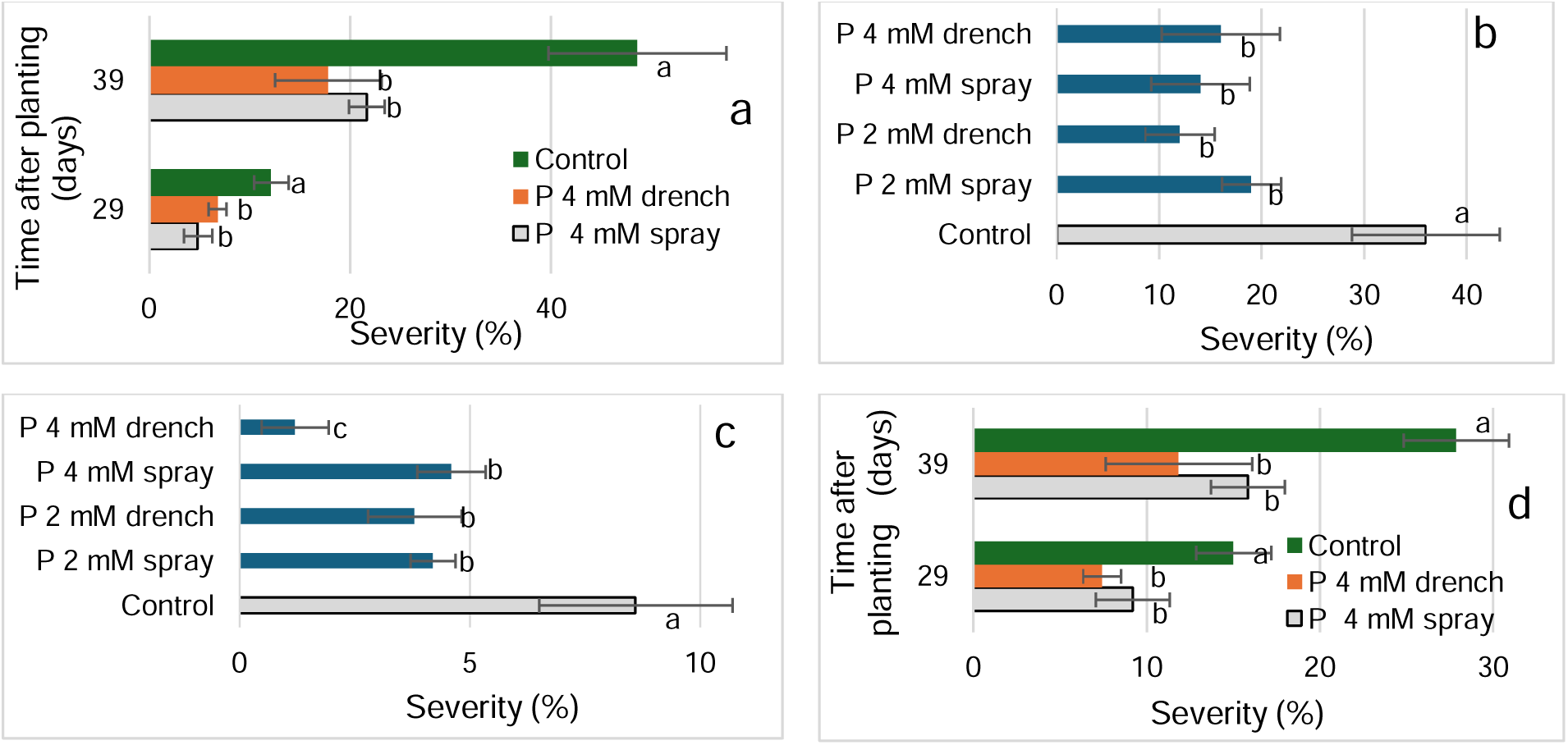
Effect of P on foliar diseases of wheat. Wheat cv. Mabruk or Bet Hashita plants were naturally infected via exposure to inoculum plants. (a) Effect of 4 mM P applied by drench or spray against *Blumeria graminis* f. sp. *tritici* (powdery mildew). Treatments were applied at 19, 26, and 33 days after seeding; disease was assessed at 29 and 39 days after treatment initiation. (b) Effect of 2- and 4-mM P (spray and drench) on *Zymoseptoria tritici* (Septoria tritici blotch). Applications were made from 14 to 67 days after seeding; disease was assessed at day 75. (c) Effect of 2- and 4-mM P (spray and drench) on *Puccinia triticina* (brown rust/leaf rust). Applications were made from 14 to 87 days after seeding; disease was assessed at day 99. (d) Effect of 4 mM P applied by drench or spray against *Puccinia striiformis* f. sp. *tritici* (stripe/yellow rust). Treatments were applied at 19, 26, and 33 days after seeding; disease was assessed at 29 and 39 days after treatment initiation. Disease severity was rated on a 0–100 scale (0 = healthy, 100 = fully symptomatic leaves). Different letters indicate significant differences (one-way ANOVA with Tukey–Kramer HSD, α = 0.05). Error bars represent SE.

*Septoria tritici* blotch (*Zymoseptoria tritici*) was reduced by 47–67% with 2–4 mM Phe applied as drench or spray (Figure 6b). Brown rust (*Puccinia triticina*) was reduced by 47–86% with 2- and 4- mM Phe applied by either method (Figure 6c). Stripe rust (*Puccinia striiformis* f. sp. *tritici*) was reduced by 58% (drench) and 55% (spray) with 4 mM Phe (Figure 6d).

### 3.6 Suppression of cucumber downy mildew (*Pseudoperonospora cubensis*) by Phe

Cucumber plants were sprayed or drenched with 4 mM Phe at 7–12 days intervals until day 50. Downy mildew developed naturally, and symptoms appeared from day 32 and were assessed at day 61 after treatment initiation (Figure 7). Disease severity reached 29% in untreated plants and was reduced by 90% (spray) and 98% (drench) with Phe (Figure 7).

**FIGURE 7.**
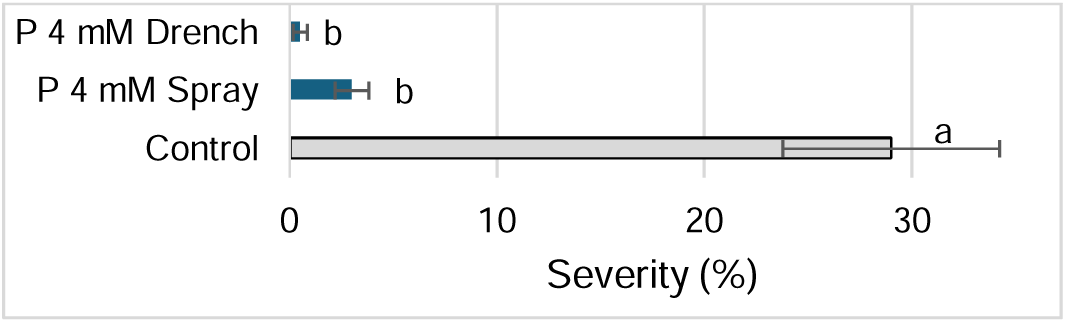
Effect of P on cucumber downy mildew (*Pseudoperonospora cubensis*). Effect of 4 mM P applied by spray or drench at 0, 7, 14, 22, 35, 42, 49, and 56 days. Downy mildew severity was assessed at day 61 on a 0–100 scale (0 = healthy, 100 = fully covered leaves). Different letters indicate significant differences (one-way ANOVA with Tukey–Kramer HSD, α = 0.05). Error bars represent SE.

### 3.7 Effect of Phe on *Pythium* damping-off in cucumber

In a short-term assay, a 4 mM Phe spray applied before transplanting into *Pythium aphanidermatum*- infested medium significantly reduced damping-off incidence (Figure S6). In additional experiments, 2- and 4-mM Phe applied before and after transplanting showed that only 4 mM Phe spray and drench significantly reduced disease, achieving 71% reduction, while the commercial oomycide Dynon drench achieved 89% reduction over 10 days (Figure 8).

**FIGURE 8.**
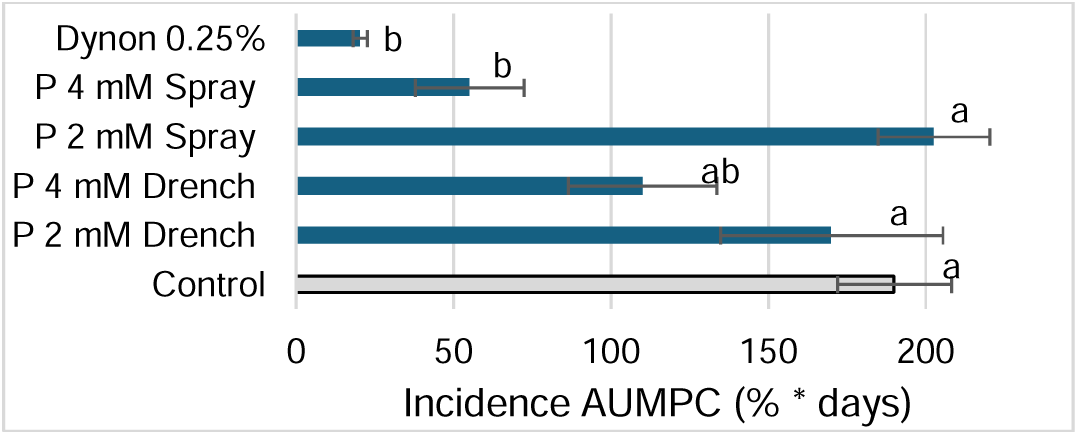
Effect of Phe on damping-off of cucumber caused by *Pythium aphanidermatum*. Seedlings were transplanted 7 days after sowing into growth medium infested with *P. aphanidermatum* (millet seed inoculum). Effect of 2- and 4-mM Phe (spray and drench) compared with Dynon 0.25% (propamocarb HCl, Bayer). Treatments were applied at 3, 6, and 10 days after sowing; mortality was monitored up to 10 days after transplanting, and AUMPC was calculated. Different letters indicate significant differences (one-way ANOVA with Tukey–Kramer HSD, α = 0.05). Error bars represent SE.

### 3.8 Growth of filamentous pathogens in culture

Phe (1–16 mM) did not inhibit linear mycelial growth in vitro (Figure S7). Instead, low concentrations stimulated growth: *B. cinerea* and *S. sclerotiorum* increased at 1–4 mM (Figure S7a,b), *R. solani* at 2 mM, and *P. aphanidermatum* at 2–16 mM (Figure S7d).

### 3.9 Effect of Phe on tomato bacterial diseases: bacterial speck (*Pseudomonas syringae* pv. *tomato*) and bacterial canker (*Clavibacter michiganensis* subsp. *michiganensis*)

Bacterial speck (PsBs) severity was reduced by 4 mM Phe by 33% (spray) and 42% (drench) (Figure 9a). Shatah 90 alone had no effect, whereas 4–12 mM Phe sprays, including 8 mM Phe combined with Shatah 90, reduced disease by 41–69% (Figure 9b). For bacterial canker (Cmm), lesion incidence was reduced by 50% with 4 mM Phe spray, while drench application had no effect (Figure 9c).

**FIGURE 9.**
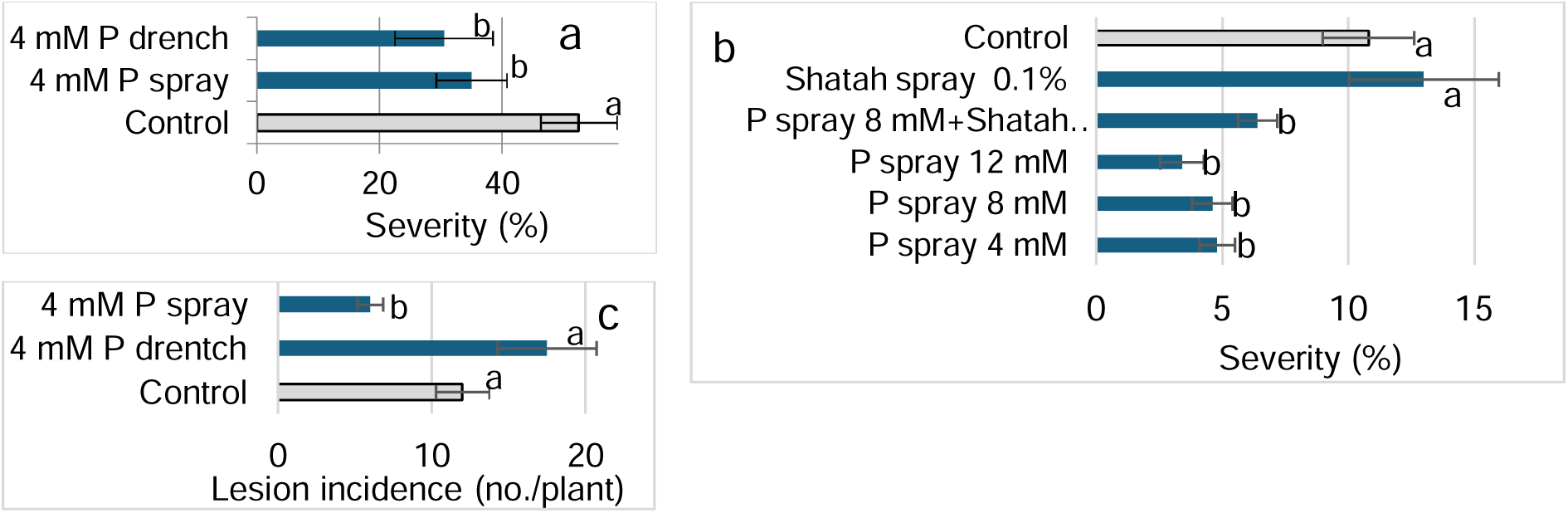
Effect of Phe on bacterial diseases of tomato: bacterial speck (*Pseudomonas syringae* pv. *tomato*) (a, b) and bacterial canker (*Clavibacter michiganensis* subsp. *michiganensis*, Cmm) (c). (a) Effect of 4 mM Phe applied by drench or spray at 0 and 4 days; disease severity assessed at day 13. (b) Effect of 4-, 8-, and 12-mM Phe sprays, Shatah 90 (alkyl phenol oxide 920 g/L, Adama/Makhteshim), and 8 mM Phe + Shatah 90 applied at days 0 and 3; disease assessed at day 14. (c) Effect of 4 mM Phe (spray and drench) applied at −3 and 0 days before Cmm inoculation; lesion incidence assessed 8 days post-infection. Bacterial speck severity was rated on a 0–100 scale (0 = no symptoms, 100 = fully covered leaves). Different letters indicate significant differences (one-way ANOVA with Tukey–Kramer HSD, α = 0.05). Error bars represent SE.

### 3.10 Effect of Phe on symptoms of tomato viral disease - Tomato brown rugose fruit virus (ToBRFV)

Symptoms of Tomato brown rugose fruit virus were followed from day 14 until day 25, reaching 36.0% and significantly reduced at all evaluation’s dates (Figure S8). The calculated AUDPC of the results in Figure S8 are presented in Figure 10. ToBRFV severity was reduced by 61 and 39% by the 4 mM Phe spray and drench, respectively (Figure 10).

**FIGURE 10.**
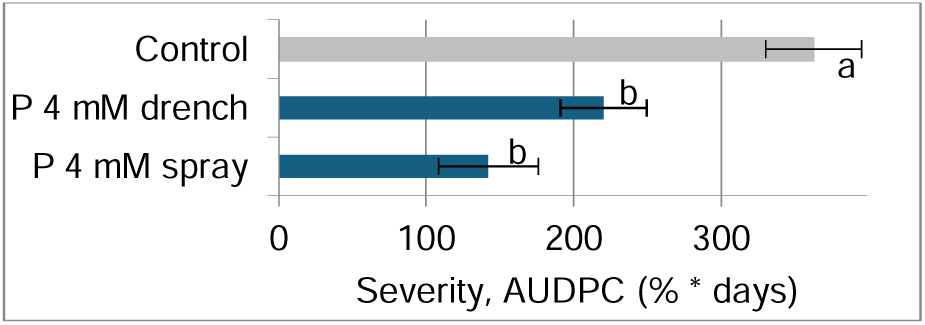
Effect of Phe on tomato viral diseases - Plants artificially infected by Tomato brown rugose fruit virus (ToBRFV) and treated by drench and spray 4 mM P, 3 times until 14 days after initiation. Symptoms severity was observed from day 14 until day 25 (a). (b.) Area under disease progress curve (AUDPC) was calculated through day 25. The severity of the virus symptoms was evaluated according to a 0 to 100% where 0= leaves with no visible symptoms and 100=leaves fully covered by virus symptoms. Values in each date followed by a different letter are significantly different according to one-way ANOVA with Tukey–Kramer’s HSD test. Default significance levels were set at α = 0.05. Bars = SEs.

The presence of the virus in leaflet samples was tested; 60% of the untreated control samples were positive for ToBRFV and it was totally reduced by the spray treatment while the drench treatment did not reduce the virus samples in leaflet samples.

## 4 Discussion

L-Phenylalanine (Phe) was recently reported to suppress *Botrytis cinerea* gray mold and several postharvest diseases. Phe is a precursor of the phenylpropanoid pathway (PPP) (Dixon et al. 2002; Oliva et al. 2020), which produces phenolics, flavonoids (flavonols and isoflavonoids), and anthocyanins. These metabolites function in plant defense signaling and resistance to pathogens, including postharvest diseases (Nicholson & Hammerschmidt 1992; Xu et al. 2018; Kumar Patel et al. 2020). We previously suggested that Phe induces a broad-spectrum defense response, as demonstrated in chrysanthemum flowers (Kumar Varun et al. 2020). Phe treatment led to the accumulation of 3-phenyllactate and benzaldehyde, along with induction of genes involved in Ca² signaling and receptor-like kinases, indicating activation of defense pathways. Phe-induced resistance was associated with priming, enabling rapid and targeted reprogramming of plant defense responses. In addition, antifungal volatiles such as phenylacetaldehyde and eugenol increased, as did coniferin, a putative monolignol precursor involved in cell wall lignification. Concurrently, ROS production and ethylene emission were reduced, while genes related to cell wall biosynthesis (including the RLK THESEUS1) and Ca² and hormonal signaling were differentially regulated (Kumar Varun et al. 2020).

Phe treatment of mango fruit induced defense-related genes involved in Ca² signaling, MAP kinase cascades, WRKY transcription factors, and activation of the phenylpropanoid pathway, leading to accumulation of flavonols and anthocyanins (Kumar Patel et al. 2023). Phenolic extracts from Phe-treated mango peel inhibited the growth of *Colletotrichum gloeosporioides*, *B. cinerea*, *Alternaria alternata*, and *Lasiodiplodia theobromae*. Overall, Phe activated fruit defense responses and stimulated the biosynthesis of antioxidant and antifungal flavonoids, effectively reducing fungal growth and disease development (Kumar Patel et al. 2023).

Similarly, Li et al. (2024) showed that Phe treatment of pear fruit suppressed *A. alternata* infection, enhanced expression and activity of phenylpropanoid-related enzymes and was accompanied by increased accumulation of caffeic acid, anthocyanins, flavonoids, lignin, and total phenolics. They further suggested that Phe enhances resistance via coordinated regulation of phenylpropanoid metabolism, MAP kinase cascades, Ca² signaling, and WRKY transcription factors. These mechanisms appear broadly conserved and effective across diverse patho-systems. Consistently, in the present study, Phe reduced disease severity in dicot crops (tomato, cucumber, sweet basil, lettuce) and in the monocot wheat. Importantly, Phe is not directly toxic to pathogens, as it does not inhibit conidial germination of *C. gloeosporioides*, *A. alternata*, *L. theobromae* (Kumar Patel et al. 2020), or *B. cinerea* (Oliva et al. 2020). Moreover, we show here that Phe does not inhibit mycelial growth of *B. cinerea*, *S. sclerotiorum*, *R. solani*, or the oomycete *P. aphanidermatum* across 1–16 mM; in some cases, pathogen growth was even stimulated at certain concentrations (Figure S7).

In tomato, *B. cinerea* was effectively controlled not only in a determinate processing cultivar typically grown in the open field (Oliva et al. 2020) but also in the greenhouse indeterminate cultivar Ikram (Figure 1 and S1). In this system, Phe drenching, was more effective than foliar spray, with a trend toward improved control at higher concentrations (Figure S1a and 1a). Phe spray performed similarly to the fungicide Switch in suppressing *B. cinerea*, while Phe drench was somewhat more effective. Phe also outperformed the fungicide Mythos; combinations showed comparable efficacy, with an additive effect observed for Mythos combined with Phe drench (Figure 1a). Except for the lowest concentration tested (2.05 mM) applied as a spray, which was ineffective on younger leaves, disease control was achieved on both young and mature leaves. Notably, efficacy extended to older leaves, where disease severity was more than twice that observed in younger leaves (Figure S1a).

We evaluated the optimal timing of Phe application prior to infection in the *B. cinerea*–tomato patho- system (Figure 1c,d). A single application was most effective when applied 5 days before infection, while combined applications at 7 and 0 days prior to infection provided superior control compared to other double-treatment schedules. Such timing effects were not assessed in other pathosystems in this study, but similar temporal dynamics have been reported for other resistance-inducing treatments.

For example, in grapevine treated with nutrient broth for control of powdery mildew (*Erysiphe necator*), defense gene expression (PR-1, PR-3, PR-6, LOX-9, OSM-1) peaked around 6 days post-treatment (Nesler et al. 2015). In tomato, root-zone warming that primed immunity was most effective against *B. cinerea* when applied 3 and 0 days before infection, and against *Xanthomonas campestris* pv. *vesicatoria* when applied 3–5 days prior to infection (Gupta et al. 2021). In contrast, in the grapevine–*Plasmopara viticola* system, *Trichoderma harzianum* T39 and BTH (bion) were most effective when applied 1 day before infection, compared with longer pre-infection intervals (Pedrazzoli et al. 2008).

Similarly to its effect on *B. cinerea* in tomato leaves, 4 mM Phe applied as either a drench or a spray effectively reduced *Sclerotinia sclerotiorum* white rot in tomato (Figure 2 and S2). Both pathogens are necrotrophs, and *S. sclerotiorum* was also suppressed in cucumber, sweet basil, and lettuce leaves. However, unlike in the *B. cinerea* system, Phe was less effective than the fungicide treatment (Figure 2b). Niklas et al. (1993) suggested that phenylalanine ammonia-lyase (PAL) contributes to partial physiological resistance of common bean to *S. sclerotiorum*. PAL catalyzes the deamination of L-phenylalanine to trans-cinnamic acid, initiating the PPP responsible for the biosynthesis of lignin, flavonoids, and other defense-related compounds (MacDonald & D’Cunha 2007).

Both necrotrophic and biotrophic pathogens were suppressed in tomato by Phe treatment (Figs. 3, S3, and 4). For *Leveillula taurica* (LtPM), higher concentrations (up to 16 mM) were effective, while 2 mM was ineffective (Figure S3). Phe was also more effective than MKP, a known powdery mildew resistance inducer (Reuveni et al. 1993; Elad et al. 2021a) (Figure 3b). Similarly, in *Oidium neolycopersici* (OnPM), higher concentrations (4–8 mM) provided better control, whereas 1 mM was ineffective (Figure S4). Amino acid–based suppression of powdery mildew has been reported previously in grapevine treated with nutrient broth, a protein hydrolysate containing amino acids (Nesler et al. 2015; Pertot & Elad 2009), but no prior reports specifically address Phe beyond our own patents (Oren-Shamir et al. 2017–2018). Here, we also show that Phe controlled *Podosphaera xanthii* (PxPM) in cucumber (Figure 5 and Figure S5). Disease suppression was evident from the cotyledon stage (Figure 5a) through mature leaves, where 2 mM was ineffective while 4 and 16 mM were effective (Figure S5a). At 4 mM, Phe outperformed MKP (Figure 5b), and drench and spray applications showed similar efficacy. Phe was comparable to Bion in controlling PxPM, although sulfur-based fungicide provided slightly stronger control (Figure 5c).

Wheat foliar diseases caused by biotrophs (*Blumeria graminis* f. sp. *tritici*—powdery mildew; *Puccinia triticina*—leaf rust; *Puccinia striiformis* f. sp. *tritici*—stripe rust) and the hemibiotroph *Zymoseptoria tritici* (septoria tritici blotch) were all effectively controlled by Phe (Figure 6), demonstrating its activity across diverse wheat pathogens. Apart from our patent (Oren-Shamir et al. 2017–2018), similar Phe-specific effects have not been reported. However, phenylalanine metabolism and the phenylalanine–tyrosine–tryptophan biosynthesis pathways were shown to contribute to maize resistance towards *Fusarium proliferatum*, a soilborne pathogen causing stalk rot (Sun et al. 2024).

Interestingly, Phe also affected oomycete- and bacteria-induced diseases in cucumber and tomato (Figs. 7, 8). In cucumber, 4 mM Phe applied as both drench and spray suppressed *Pseudoperonospora cubensis* (downy mildew) (Figure 7), while only 4 mM spray effectively controlled the soilborne damping-off pathogen *Pythium aphanidermatum*, performing similarly to the pesticide Dynon (Figure 8). Lower concentrations (2 mM spray and 2–4 mM drench) were ineffective against *P. aphanidermatum*, likely due to insufficient systemic uptake to the shoots (Figure 8).

In tomato, Phe reduced diseases caused by the Gram-negative bacterium *Pseudomonas syringae* pv. *tomato* (bacterial speck) and the Gram-positive *Clavibacter michiganensis* subsp. *michiganensis* (bacterial canker). In both cases, ≥4 mM foliar spray was effective, while drench application was effective only against bacterial speck (Figure 9).

These findings align with reports linking phenylalanine ammonia-lyase (PAL) activity to enhanced disease resistance. PAL correlates with increased phenolic content and resistance in tomato against *C. michiganensis* (Umesha, 2006) and contributes to resistance against *Ralstonia solanacearum* together with polyphenol oxidase (Vanitha et al. 2009). Similarly, PAL-dependent phenylpropanoid activation was implicated in resistance to *Pythium myriotylum* in ginger (Augustine et al. 2024), supporting the idea that targeting this pathway may improve disease management. In cucumber, induced systemic resistance against *P. cubensis* was associated with elevated defense enzymes (peroxidase, polyphenol oxidase, PAL, β-1,3-glucanase, chitinase, catalase) and increased phenolic compounds (Anand et al. 2007).

Interestingly, Phe applied to tomato plants decreased the severity of virus disease symptoms as was demonstrated for ToBRFV (Figure 10). There are very few indications of induction of resistance towards ToBRFV. Zohoursoleimani et al (2025) Reported two bacterial isolates (*Pseudomonas fluorescens* and *Burkholderia gladioli*) that induce resistance against the virus.

We evaluated several approaches to enhance the disease control efficacy of Phe. Increasing its concentration improved control of *B. cinerea* in tomato (Figure S1a), *Sclerotinia sclerotiorum* in lettuce (Figure 2c), and the powdery mildews *Leveillula taurica* (Figure S3) and *Oidium neolycopersici* (Figure S4) in tomato. However, higher concentrations did not improve control of *Podosphaera xanthii* in cucumber (Figure S5a) or *Pseudomonas syringae* pv. *tomato* in tomato (Figure 9b).

Co-application with an adjuvant improved performance in both tomato powdery mildew systems, where the combination showed synergistic effects compared with either component alone (Figs. 3d, 4b). In contrast, the adjuvant did not enhance activity against *P. syringae* pv. *tomato* (Figure 9b).

Increasing application frequency (splitting weekly 4 mM treatments into two 2 mM sprays) did not improve control in either *L. taurica*–tomato or *P. xanthii*–cucumber systems (Figs. 3b, S5b), consistent with reports where application frequency can influence biocontrol efficacy (Shafir et al. 2006).

Formulation of control agents may prove to be more effective than the non-formulated product (Hazra & Purkait 2019). Indeed, here a commercial formulation containing 55% active ingredient (NaturaFend 550 SP; ICA International Chemicals) enhanced disease control compared with non-formulated Phe, improving suppression of *B. cinerea* in tomato (Figure 1d) and *P. xanthii* in cucumber (Figure 5c). This product is currently registered in South Africa for control of postharvest and field diseases including anthracnose (*Colletotrichum gloeosporioides*) in avocado and mango, botrytis bunch rot and powdery mildew in grape, and stem-end rot in mango (*Botryosphaeria* spp.).

As shown throughout the results section, Phe was effective as both foliar spray and a drench against foliar diseases. The root-to-canopy activity observed following drench application may result from the translocation of phenylalanine and/or its conversion into phenylpropanoid-derived metabolites, which function as defense signaling molecules in leaves (Nicholson & Hammerschmidt 1992; Xu et al. 2018). Alternatively, defense-related compounds may be synthesized in roots and transported systemically to the canopy. In addition, phenylalanine feeds into the PAL pathway for salicylic acid biosynthesis in plants (Zhu et al. 2025). Salicylic acid is a key phytohormone that regulates responses to biotic stress (Gupta et al. 2021; Perazzolli et al. 2009), and it likely contributes to Phe-induced resistance towards pathogens.

In conclusion, phenylalanine applications, as either drench or spray, effectively controlled foliar and soilborne diseases across a wide range of patho-systems, including fungal, oomycete, bacterial and viral pathogens that have necrotrophic, biotrophic, and hemi-biotrophic lifestyles in both dicot and monocot species. Our recent data (Elad and Oren-Shamir, unpublished results) also prove that Phe is effective in cucumber and tomato against the viruses Cucurbit yellow stunting disorder virus (CYSDV), Cucurbit chlorotic yellows virus (CCYV) in cucumber, and Tomato yellow leaf curl virus (TYLCV) that are transmitted by *Bemisia tabaci* in cucumber and tomato, the arthropod pests *B. tabaci*, *Tetranychus urticae*, *Liriomyza bryoniae* in tomato and cucumber, *L. trifolii* and *Spodoptera littoralis* in lettuce. and *Tuta absoluta* in tomato. This broad activity suggests diverse and general mechanisms of action. Effective control was achieved at relatively low concentrations (≥4 mM), with improved performance observed at higher doses and further enhancement when combined with an adjuvant or applied as the formulated product NaturaFend.

## ACKNOWLEDGEMENTS

We thank the late Eyal Cohen RIP, Dr. Galit Sharabani, and Ohad Zuckerman of Copia for the encouragement and advice during the PostBoost project development and for the support. We thank members of the Elad and Oren-Shamir research groups at the Volcani Institute for their continuous support. The fruitful discussions and help of Dr. Noam Alkan of the Institute of Postharvest and Food Sciences, Volcani Institute is much appreciated. We thank the following people of the Plant Pathology Department: Dr. Omer Frenkel and Gidon Mordocovitz for support, advice and supply of *Pythium* and *Rhizoctonia* inocula, Dr. Aviv Dombrovski for the ToBRFV inoculation and presence evaluation, Dr. Doron Tepper for the advice regarding bacterial diseases and cmm inoculum, and Menachem Borenshtein, Eyal Glanz, and Ran Shulhani for the technical support in the greenhouses. The warm and interesting discussions and the excellent implementation of the following people of ICA International Chemicals (Pty) Ltd. South Africa is very much appreciated: Wouter Schreuder Jr., the late Kobus Serfontein RIP, Suzel Serfontein, Pieter Louw, and Gert van Coller. We thank Arik Hirshman formerly in Agridera Seed &. Agriculture Ltd, and currently in Hazera 1939 Ltd for the wheat Mabruk seeds and diseased plants.

## Author Contributions

**Yigal Elad**: Writing – review & editing, Writing – original draft, Supervision, Methodology, Investigation, Formal analysis, Conceptualization. **Michal Oren Shamir**: Methodology, writing – review & editing, Supervision, Methodology, Investigation, Funding acquisition, Formal analysis, Conceptualization. **Dalia Rav-David**: Methodology, Investigation, Formal analysis.

## Data Availability Statement

The authors declare that the data supporting the findings of this study are available within the paper and its Supporting Information files. Raw data is available from the corresponding author upon reasonable request.

