## Supplemental material for "Phenylalanine Treatment Suppresses Necrotrophic and Biotrophic Pathogens in Mono- and Dicotyledonous Plants"

Supplemental materials

a

Disease severity, AUDPC (%*days)

| Leaves | 0 | P 2.05 mM Spray | P 4.10 mM Spray | P 8.20 mM Spray | P 2.05 mM Drench | P 4.10 mM Drench | P 8.20 mM Drench |
| --- | --- | --- | --- | --- | --- | --- | --- |
| Middle | 223.062 | 170.25 | 151 | 129.3 | 97.25 | 94.65 | 57.55 |
| Upper | 100.45 | 83.175 | 54.525 | 54.675 | 22.075 | 19.625 | 9.025 |

b

Disease severity, AUDPC (%*days)

|  | AUDPC |
| --- | --- |
| Control | 126.351 |
| Switch 0.1% | 31.1197 |
| P 1.0 mM Spray | 27.225 |
| P 2.05 mM Spray | 25.05 |
| P 4.1 mM Spray | 28.2 |
| P 1.0 mM Drench | 31.725 |
| P 2.05 mM Dench | 2.475 |
| P 4.1 mM Drench | 1.5 |

**FIGURE S1**

Control of *Botrytis cinerea* gray mold (BcGM) on tomato cv. Ikram by Phe and the fungicide switch.

(a) Effect of leaf age and Phe concentrations (2.05–8.20 mM) applied by spray or drench on BcGM. Treatments were applied at −3 and 0 days before *B. cinerea* conidia infection and assessed up to 12 days post-infection. (b) Effect of Phe (1.0–4.1 mM) applied by spray or drench compared with Switch (cyprodinil + fludioxonil, Syngenta). Treatments were applied at −3 and 0 days before infection and evaluated up to 11 days post-infection. BcGM severity after conidial infection was rated on a 0–100% scale (0 = healthy; 100 = maximum rot). After mycelial disc infection, severity reflects lesion area (e, f). Area under the disease progress curve (AUDPC) was calculated over the assessment period. Different letters indicate significant differences (one-way ANOVA with Tukey–Kramer HSD, α = 0.05). Error bars represent SE.

Severity, AUDPC (mm2*days)

|  | AUDPC |
| --- | --- |
| Control | 175.375 |
| P 4 mM spray | 60.375 |
| P 4 mM drench | 102.25 |

**FIGURE S2**

Effect of P on *Sclerotinia sclerotiorum* white rot (SsWM) caused by mycelial disc infection on sweet basil, cucumber, and lettuce leaves. Sweet basil SsWM control by 4 mM P spray or drench applied at −4 and 0 days before inoculation; disease assessed over 6 days. Disease severity was quantified by lesion diameter and converted to rot area. Area under the disease progress curve (AUDPC) was calculated. Different letters indicate significant differences (one-way ANOVA with Tukey–Kramer HSD, α = 0.05). Error bars represent SE.

a

Severity (%)

|  | 24 |
| --- | --- |
| 0 | 5.4 |
| 2 mM spray | 3.6 |
| 2 mM drench | 4.3 |
| 4 mM spray | 1.5 |
| 4 mM drench | 2.9 |
| 16 mM spray | 0.833333 |
| 16 mM drench | 1.35 |

**FIGURE S3**

Effect of P on *Leveillula taurica* powdery mildew (LtPM) of tomato. Effect of P applied by spray or drench at 2–16 mM. Treatments were applied at −4 and 0 days before inoculation and at 7- and 14-days post-inoculation; disease severity was assessed at day 24. Disease severity was rated on a 0–100 scale (0 = healthy, 100 = fully covered leaves). Different letters indicate significant differences (one-way ANOVA with Tukey–Kramer HSD, α = 0.05). Error bars represent SE.

Severity (% coverage)

|  | 46 |
| --- | --- |
| 0 | 20.4 |
| Spray 1 mM | 4.2 |
| Spray 2 mM | 3.6 |
| Spray 4 mM | 2 |
| Spray 8 mM | 0.6 |
| Drench 1 mM | 24.4 |
| Drench 2 mM | 7.6 |
| Drench 4 mM | 0 |
| Drench 8 mM | 0 |

**FIGURE S4**

Effect of P on *Oidium neolycopersici* powdery mildew (OnPM) of tomato. Effect of P applied by spray or drench at 2–8 mM. Treatments were applied at −4 and 0 days before inoculation and then weekly until 35 days post-inoculation; disease severity was assessed at day 46. Disease severity was rated on a 0–100 scale (0 = healthy, 100 = fully covered leaves). Different letters indicate significant differences (one-way ANOVA with Tukey–Kramer HSD, α = 0.05). Error bars represent SE.

a

Severity, AUDPC (% * days)

|  | AUDPC 29 d |
| --- | --- |
| 0 | 60.275 |
| 2 mM spray | 40.875 |
| 4 mM spray | 26.175 |
| 2 mM drench | 44.775 |
| 4 mM drench | 21.0625 |
| 16 mM drench | 29.4375 |

b

Severity (%)

|  | 50 |
| --- | --- |
| Control | 14 |
| 2 mM once/week | 14.4 |
| 4 mM once/week | 8.4 |
| 2 mM twice/week | 11.8 |

**FIGURE S5**

Effect of P on *Podosphaera xanthii* powdery mildew (PxPM) of cucumber. (a) Effect of P applied at −4 and 0 days before inoculation and three times weekly for 3 weeks; disease severity was assessed at day 29. (b) Effect of 2 mM P applied once or twice weekly and 4 mM P applied once weekly for 6 weeks; disease was assessed at 50 days after treatment initiation. Disease severity was rated on a 0–100 scale (0 = healthy, 100 = fully covered leaves). AUDPC was calculated (a). Different letters indicate significant differences (one-way ANOVA with Tukey–Kramer HSD, α = 0.05). Error bars represent SE.

Dead plants (%)

|  | % dead plants |
| --- | --- |
| Untreated | 43.75 |
| P 4 mM | 6.25 |

**FIGURE S6**

Effect of Phe on damping-off of cucumber caused by *Pythium aphanidermatum*. Seedlings were transplanted 7 days after sowing into growth medium infested with *P. aphanidermatum* (millet seed inoculum). Effect of 4 mM Phe spray applied before transplanting. Seedlings were transplanted 4 days after treatment and mortality was assessed 3 days later. Different letters indicate significant differences (one-way ANOVA with Tukey–Kramer HSD, α = 0.05). Error bars represent SE.

a

Growth rate (cm2/day)

| Phenylalanine concentration (mM) | days 3to5 |
| --- | --- |
| 0 | 13.2469 |
| 1 | 15.7968 |
| 2 | 16.6799 |
| 4 | 16.6799 |
| 8 | 13.4431 |
| 16 | 13.7611 |

b

Growth rate (cm2/day)

| Phenylalanine concentration (mM) | Sclerotinia days 2 to 4 |
| --- | --- |
| 0 | 13.5412 |
| 1 | 14.5251 |
| 2 | 14.5408 |
| 4 | 14.4396 |
| 8 | 13.3938 |
| 16 | 13.5822 |

c

Growth rate (cm2/day)

| Phenylalanine concentration (mM) | Days one to three |
| --- | --- |
| 0 | 17.1326 |
| 1 | 17.9189 |
| 2 | 19.7977 |
| 4 | 18.4998 |
| 8 | 18.0184 |
| 16 | 16.3699 |

d

Growth rate (cm2/day)

| Phenylalanine concentration (mM) | 2 |
| --- | --- |
| 0 | 20.1195 |
| 1 | 21.8204 |
| 2 | 23.7332 |
| 4 | 25.0389 |
| 8 | 24.4946 |
| 16 | 27.0485 |

**FIGURE S7**

Effect of P (1–16 mM) incorporated into growth medium on mycelial growth of *Botrytis cinerea* (a), *Sclerotinia sclerotiorum* (b), *Rhizoctonia solani* (c), and *Pythium aphanidermatum* (d). Colony diameter was measured 1–4 days after disc inoculation, and growth rate per day was calculated over 3–5 days (a), 2–4 days (b), 1–3 days (c), and 1–2 days (d), depending on pathogen. Error bars represent SE at each P concentration.

a

Intensity (% affected leaflets)

| Time after treatment (days) | 4 mM P spray | 4 mM P drench | Control |
| --- | --- | --- | --- |
| 11 | 0 | 0 | 0 |
| 14 | 4 | 11 | 20 |
| 15 | 6 | 13 | 23 |
| 21 | 15 | 18 | 28 |
| 25 | 16 | 20 | 36 |

**FIGURE S8**

Effect of Phe on tomato viral diseases - plants artificially infected by Tomato brown rugose fruit virus (ToBRFV) and treated by drench and spray 4 mM P, 3 times until 14 days after initiation. Symptoms severity was observed from day 14 until day 25. Area under disease progress curve (AUDPC) was calculated through day 25 and is presented in Fig. 10. The severity of the virus symptoms was evaluated according to a 0 to 100% where 0= leaves with no visible symptoms and 100=leaves fully covered by virus symptoms. Values in each date followed by a different letter are significantly different according to one-way ANOVA with Tukey–Kramer’s HSD test. Default significance levels were set at α = 0.05. Bars = SEs.
